# Single Cell Transcriptomics Reveal Genomic Indicators of Nonalcoholic Fatty Liver Disease

**DOI:** 10.1101/2024.11.04.621970

**Authors:** Madhavendra Thakur

## Abstract

Nonalcoholic fatty liver disease (NAFLD) affects roughly 25% of people globally. This paper seeks to use single cell RNA sequencing (scRNA-seq) to identify genomic markers for the identification of NAFLD and the evaluation of therapeutic methods *in silico*. Publicly available scRNA-seq datasets of livers of both control and NAFLD mice were analyzed in R using the Seurat package for scRNA-seq analysis. Datasets were clustered as needed using the Seurat standard preprocessing workflow, and the zonation of hepatocyte clusters was identified based on established marker genes. When examining hepatocyte reprogramming during NAFLD, trends were seen in the zonation of hepatocytes expressing fructose metabolism pathways such as Aldob, Slc2a5 and Khk. In one dataset, healthy livers showed elevated expression of fructose metabolism pathways in central hepatocytes, while NAFLD livers exhibited this expression predominantly in portal hepatocytes. Conversely, in a second dataset, this zonation pattern was reversed, with central hepatocytes in NAFLD livers showing higher fructose metabolism expression compared to portal regions. This consistent observation of a zonation switch across datasets strengthens our understanding of hepatocyte reprogramming during NAFLD and fibrosis. Further, the direction of this switching may provide insights into fibrosis etiology, potentially reflecting differences in diet and metabolic stressors that contribute to fibrosis progression. It could also prove valuable in *in silico* evaluations of therapies for NAFLD by identifying liver fibrosis.

## INTRODUCTION

Nonalcoholic fatty liver disease (NAFLD) affects 100 million Americans (1). According to the NIH, NAFLD is a disease caused by excess fat deposits in the liver and is linked to health conditions such as obesity or diabetes (1). It can also take the form of nonalcoholic steatohepatitis (NASH), a form of NAFLD which can cause scarring of the liver, or fibrosis (1).

Single cell RNA sequencing (scRNA-seq) is a novel technology which allows for analysis of gene profiles of individual cells, thus allowing for granular understanding of gene expression profiles. Recently, scRNA-seq has been used to analyze novel therapies for NAFLD and better understand the liver microbiome, particularly with regards to non-parenchymal cells. Existing work has established that expression of fructose metabolism pathways in hepatic stellate cells (HSCs) can contribute to fibrosis (2).

This paper sought to identify if hepatocytes also displayed genomic identifiers of liver fibrosis. Particularly, it sought to do so by leveraging scRNA-seq technology to gain insight into hepatocyte subgroups. A genomic indicator was found in the zonation of hepatocytes expressing fructose metabolism genes and was confirmed when working with another data set. This could allow for fully *in silico* identifications of potential therapies for liver fibrosis and serve as a tool for further analysis of scRNA-seq datasets of the liver microbiome.

## RESULTS

Clustering analysis having grouped hepatocytes according to zonation, and marker genes having been used to identify the zonation of said cluster, the data was ready to be analyzed, and trends in the expression of genes related to fructose metabolism and fructose transport pathways, such as Khk and Slc2a5 respectively, could be observed.

The results obtained by the dataset provided by Wang, et al. can be seen in **Figure 4**. In healthy livers, Khk and Slc2a5 were expressed largely in periportal hepatocytes, while pericentral hepatocytes did not show significant levels of Khk and Slc2a5 expression. In NASH livers however, Khk and Slc2a5 were expressed largely in pericentral hepatocytes, while pericentral hepatocytes did not show significant levels of Khk and Slc2a5 expression. There is a marked switch in the zonation of hepatocytes expressing Khk and Slc2a5 between control and NASH diets.

A similar trend can be found in the data given by Coassolo et al. **Figure 5** shows the same switch in zonation of hepatocytes expressing fructose metabolism pathways. However, the switch goes in the opposite direction: expression of fructose metabolism pathways is elevated in central/midlobular areas of mice fed a control diet, and elevated in portal areas of mice fed a NAFLD diet. The results obtained by the first dataset are shown in Figure 4.

Hepatocytes display an increase in expression of fructose metabolism in central areas in NASH livers, while displaying higher expression levels of the same genes in portal areas in control mice.

Thus, the data displayed a pronounced trend in the zonation of hepatocytes expressing fructose metabolism pathways.

## DISCUSSION

The increase in expression of fructose metabolism pathways in portal areas falls in line with current understanding of the zonation of fructose metabolism (3). Specifically, increased expression of metabolic pathways in central areas represents lipogenesis, contributing to fatty liver (3). The patterns found in the dataset provided by Coassolo, 2023 demonstrate a similar reprogramming of fructose metabolism pathways, although with the inverse effect.

As the switching mechanism seen in both datasets emerges as a valuable metric for identifying fibrosis, the different directions of that switch pose further questions as to the origin of the discrepancy. One source could be the exact mice models used in each experiment. While Wang, 2023 used a “FAT-NASH” model which fed the experimental group a high fat, high fructose diet for 12 weeks, the mice in Coassolo, 2023 were fed a NASH diet for 9 weeks (4)(5). Additionally, whereas in Wang, 2023, the mice were fed a “Western diet” of 21.2% fat, 41% sucrose, and 1.25% cholesterol, in Coassolo, 2023, the mice were fed a “NAFLD diet” of 40% fat, 20% kcal fructose, and 2% cholesterol (4)(5). That reversal of levels of fat and cholesterol could be the reason the zonation of fructose metabolism differed. Indeed, this would align with existing understandings of zonation of metabolic pathways expression, where increased expression in portal areas aligns with gluconeogenesis, which could occur if the body has an excess of fats (3). If so, the direction of switching could further be used as a metric to understand the cause of liver fibrosis *in silico*.

Regardless, it can be observed that the switching in both graphs is very strongly related to fibrosis and corroborates with existing literature. This offers a useful genomic identifier for fibrosis. Despite the discrepancies in direction of switching, the granular insight that scRNA-seq offers into zonation of fructose metabolism in hepatocytes could prove valuable in future evaluation of possible therapies against fibrosis. When evaluating therapies, the zonation of fructose metabolism in hepatocytes of positive and negative controls could be used as benchmarks against which experimental sets could be compared. The different directions of hepatocyte reprogramming during NAFLD can further be used to gain insight into the dietary causes of NAFLD in particular samples *in silico*, and potentially elucidate the mechanisms driving NAFLD more broadly.

## MATERIALS AND METHODS

All programs used in the processing of data can be found in this GitHub repository: https://github.com/mThkTrn/nafld_metabolism_zonation_analysis.

Due to the rise of scRNA-seq as a tool for analysis of NAFLD, there are many public datasets available to analyze. Datasets with mice fed control diets as well as high fat, high fructose diets which induce NAFLD and fibrosis were downloaded from the National Center for Biotechnology Information’s Gene Expression Omnibus (GEO). Wang, et al. were contacted to obtain the preprocessed Seurat object used in their paper. 2 datasets, generated as part of 2 different experiments, were considered to hedge experimental bias.

The Seurat package for analysis of scRNA-seq data in the R programming language was used. Code was written in R markdown files in the RStudio editor (6). The first dataset was available preprocessed following the Seurat preprocessing workflow, and thus was immediately prepared for analysis (3).

The second dataset was downloaded from GEO as matrices of barcodes and features (5). These were then integrated into a Seurat object using the Seurat standard preprocessing workflow. Clustering analysis was performed locally, and the optimal cluster size was determined following the Seurat preprocessing workflow and steps indicated in the paper regarding the original dataset. Specifically, clustering was done to maximize variability in the set of marker genes outlined in the original paper, although the exact same clusters were not able to be reproduced (4). 15 clusters were identified. Marker genes for hepatocytes provided in the original paper were then used to identify hepatocyte clusters, as shown in Figure 1 (5).

**Figure 1.**
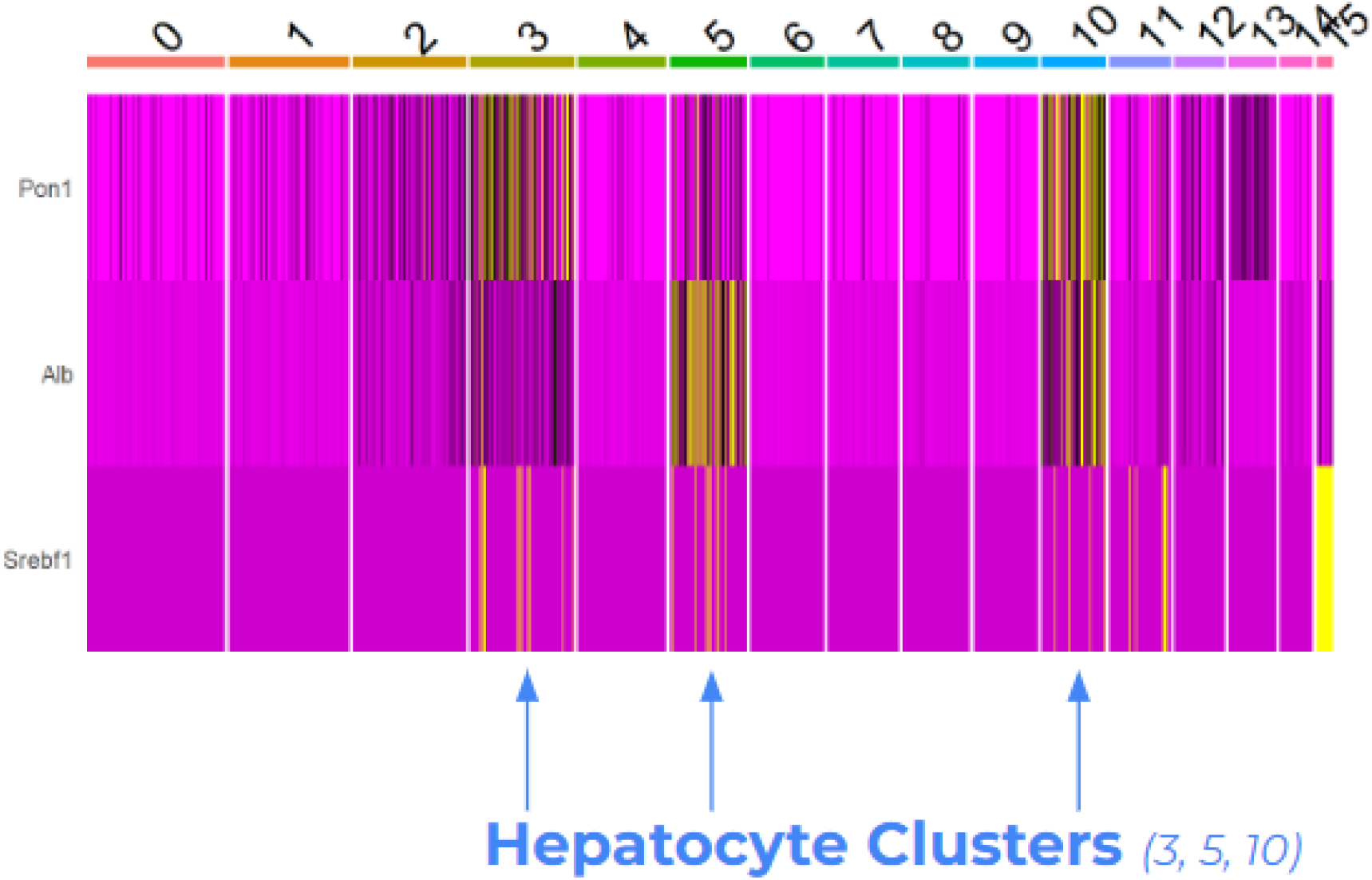
Heatmap of expression of hepatocyte marker genes in clusters, data from Coassolo, et al. Expression of Pon1, Alb and Srebf1 is shows across clusters 0 to 15 (yellow indicates expression in one cell of the cluster). Generated using Seurat in RStudio.

As clusters 3, 5 and 10 were established as hepatocyte clusters, their zonation was established using common hepatocyte marker genes for zonation (7). Hepatocytes generally fall into 3 main categories in terms of their place in the liver: pericentral, periportal or midlobular, classified by proximity to the portal and central veins, or being roughly in between the 2 respectively. Figure 2 demonstrates the use of a heatmap for classification of zonation.

**Figure 2.**
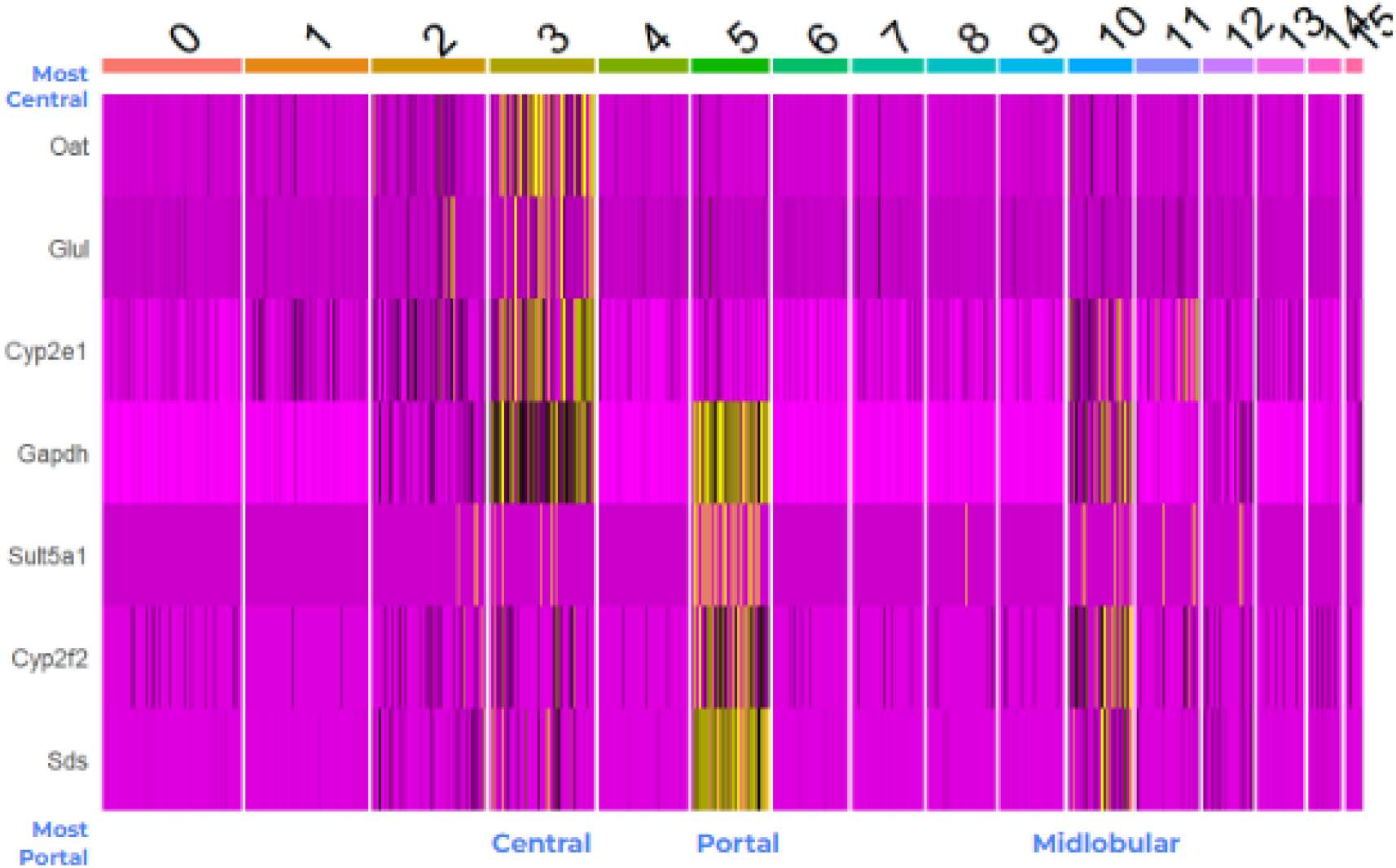
Heatmap of clusters by expression of common hepatocyte zonation markers, from most central to most portal, data from Coassolo, et al. Heatmap of expression of hepatocyte zonation markers (Oat, Glul, Cyp2e1, Gapdh, Sult5a1, Cyp2f2, Sds) across clusters 0 to 15 (yellow indicates expression in one cell of the cluster). Generated using Seurat in RStudio.

Similar analysis was done with the first dataset to establish the zonation of hepatocyte clusters, as seen in Figure 3.

**Figure 3.**
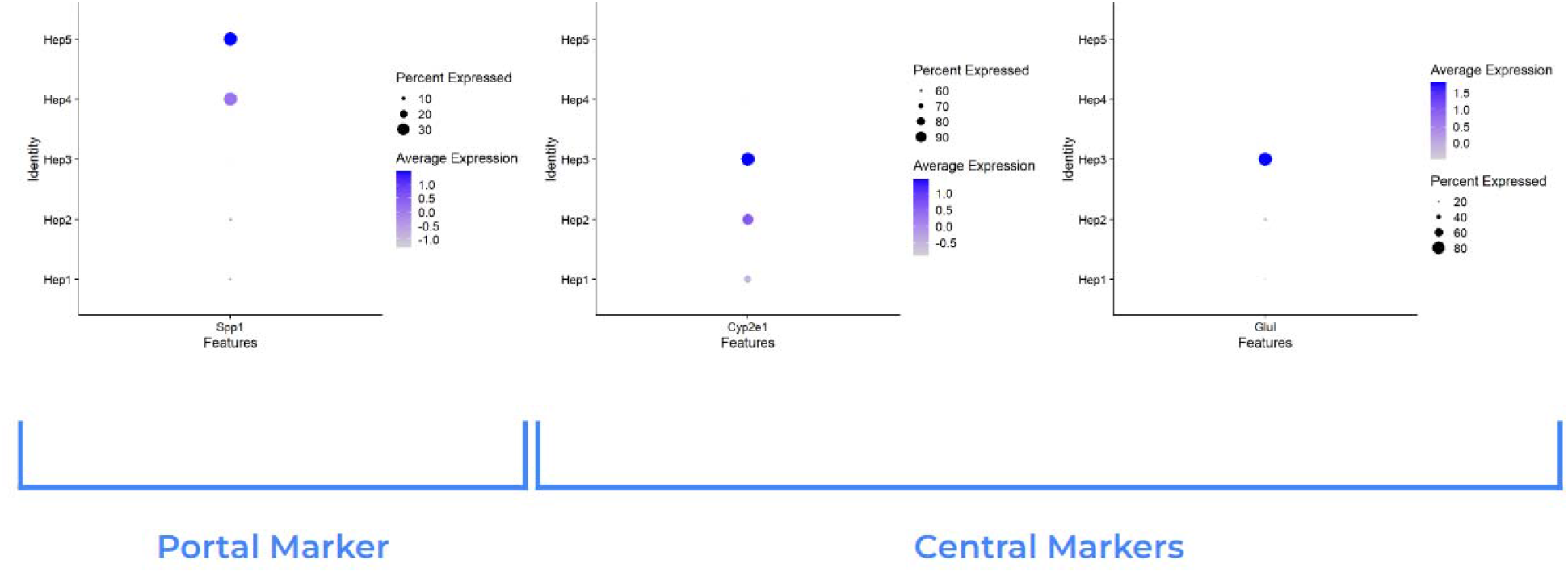
Dot plot of expression hepatocyte zonation markers across hepatocyte clusters, data from Wang, et al. 3 dot plots of expression of Spp1, Cyp2e1 and Glul respectively across all clusters. Generated using Seurat in RStudio.

**Figure 4.**
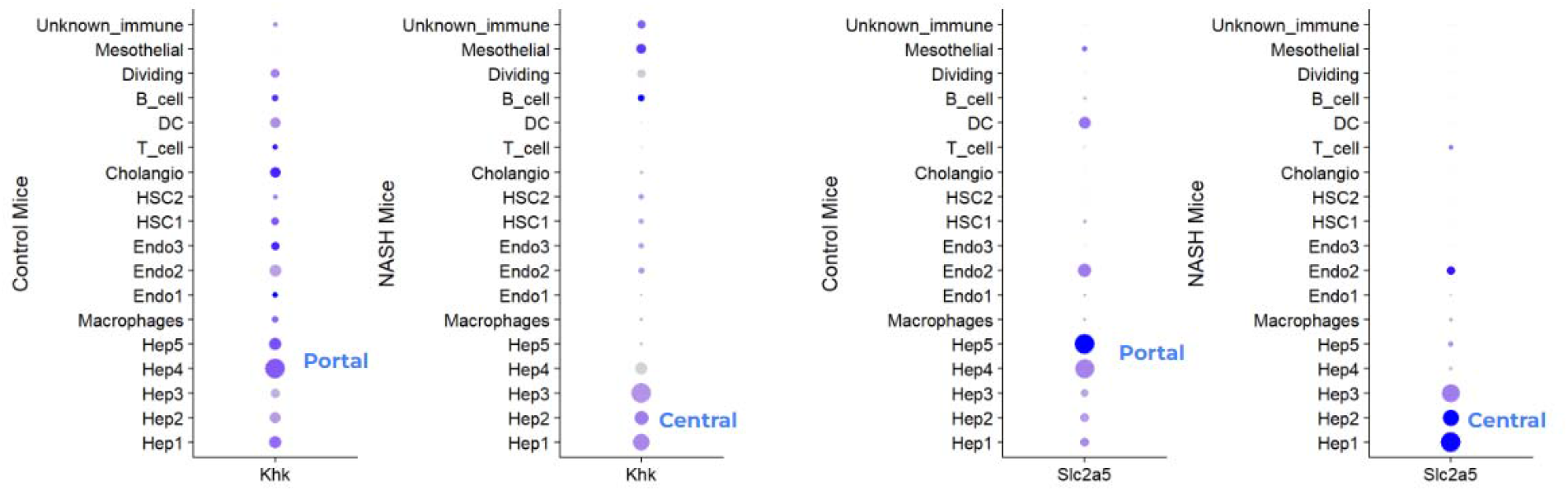
Dot plot of expression of Khk and Slc2a5 in control mice and mice fed NASH diets, data from Wang et al. 2 dot plots (left) showing expression of Khk in control and NASH mice respectively across all clusters, and 2 dot plots (right) showing expression of Slc2a5 in control and NASH mice respectively across all clusters (large dots signify increased expression relative to other clusters, darker blues signify increased average expression within clusters). Generated using Seurat in RStudio

**Figure 5.**
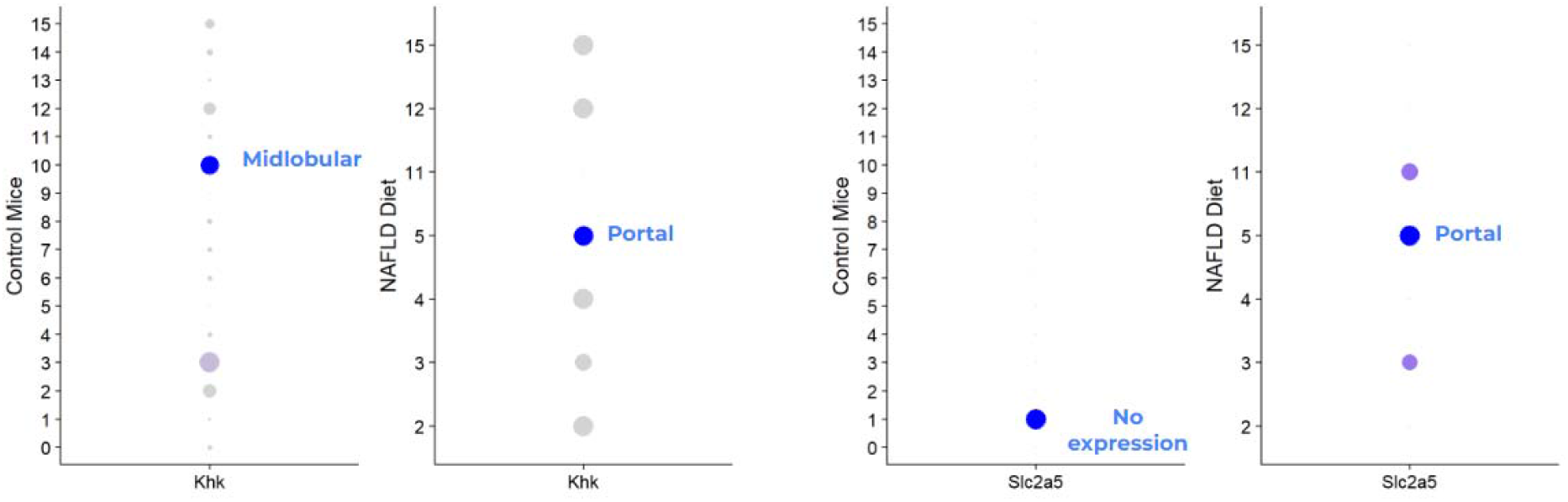
Dot plot of expression of Khk and Slc2a5 in control mice and mice fed NASH diets, data from Coassolo et al. 2 dot plots (left) showing expression of Khk in control and NASH mice respectively across all clusters, and 2 dot plots (right) showing expression of Slc2a5 in control and NASH mice respectively across all clusters (large dots signify increased expression relative to other clusters, darker colors signify increased average expression within clusters). Generated using Seurat in RStudio

Thus, with hepatocyte clusters zonated and identified in both datasets, the datasets were ready for analysis. Gene expression dot plots were generated with Seurat, plotting the expression of genes which correlate to fructose metabolism pathways. In particular, *Khk, Aldob, Slc2a5* and *Slc2a2* were analyzed, as they play roles in the transport and metabolism of fructose, and NAFLD diets are characterized by high fructose intake.

## ACKNOWLEDGMENTS (Optional)

Thank you to the Chu lab at the Icahn School of Medicine at Mount Sinai for their support of this project.

## REFERENCES

(1) Teng, Margaret LP et al. “Global incidence and prevalence of nonalcoholic fatty liver disease.” Clinical and molecular hepatology vol. 29, Suppl (2023): S32–S42. doi:10.3350/cmh.2022.0365

(2) Hou, Wei, and Wing-Kin Syn. “Role of Metabolism in Hepatic Stellate Cell Activation and Fibrogenesis.” Frontiers in cell and developmental biology vol. 6 150. 12 Nov. 2018, doi:10.3389/fcell.2018.00150

(3) RP, Porat-Shliom N. Liver Zonation – Revisiting Old Questions With New Technologies. Frontiers. 2021;12

(4) Wang S, Li K, Pickholz E, Dobie R et al. An autocrine signaling circuit in hepatic stellate cells underlies advanced fibrosis in nonalcoholic steatohepatitis. Sci Transl Med 2023 Jan 4;15(677)

(5) Coassolo L, Liu T, Jung Y, Taylor NP, Zhao M, Charville GW, et al. Mapping transcriptional heterogeneity and metabolic networks in fatty livers at single-cell resolution. iScience [Internet]. 2023 Jan 20;26(1).

(6) Hao, Y., Stuart, T., Kowalski, M.H. et al. Dictionary learning for integrative, multimodal and scalable single-cell analysis. Nat Biotechnol (2023). 10.1038/s41587-023-01767-y

(7) Umbaugh DS, Ramachandran A, Jaeschke H. Spatial Reconstruction of the Early Hepatic Transcriptomic Landscape After an Acetaminophen Overdose Using Single-Cell RNA-Sequencing. Toxicol Sci. 2021 Aug 3;182(2):327–345. doi: 10.1093/toxsci/kfab052. PMID: 33983442; PMCID: PMC8331154.

